# Increased synaptic facilitation and exploratory behavior in mice lacking the presynaptic protein Mover

**DOI:** 10.1101/560896

**Authors:** Julio S. Viotti, Frederik W. Ott, Eva M. Schleicher, Jannek M. Wagner, Yvonne Bouter, Thomas A. Bayer, Thomas Dresbach

## Abstract

In vertebrates and invertebrates, neurotransmitter release relies on a highly conserved molecular machinery. A surprisingly small number of presynaptic proteins evolved specifically in vertebrates. How they expand the power or versatility of the conserved core machinery is unclear. One of these vertebrate-specific proteins, called Mover / TPRGL / SVAP30, is heterogeneously expressed throughout the brain, suggesting that it adds special functions to subtypes of presynaptic terminals. In this study we generated Mover knockout mice to investigate the role of Mover from synaptic transmission to behavior. Deletion of Mover did not affect synaptic transmission at CA3 to CA1 synapses. In contrast, Mover deficient mice had strongly increased short-term facilitation at mossy fiber to CA3 synapses. This increase included frequency facilitation, a hallmark of mossy fiber terminal function. The effect was age- and Ca^2+^-dependent, and relied on the Kainate receptor/cAMP pathway in the mossy fiber terminals. Despite this change in presynaptic plasticity, the absence of Mover did not affect long-term spatial reference memory or working memory, but led to reduced anxiety. These discoveries suggest that Mover has distinct roles at different synapses. At mossy-fiber terminals, it acts to constrain the extent of presynaptic facilitation. Its role in activity-dependent neurotransmission could be necessary for normal anxiety responses.

**Significance Statement:** The enormous increase in the complexity of brains during evolution is accompanied by a remarkably small number of new, vertebrate-specific presynaptic proteins. These proteins are unlikely to be essential for transmitter release, because invertebrate synapses do not need them. But what functions do they fulfill? We show that the vertebrate-specific protein Mover is involved in constraining the release of neurotransmitters in some synapses in the hippocampus, while not affecting others. We further demonstrate that the absence of this protein leads to decreased anxiety levels. Understanding the function of such a protein can help us further understand synaptic transmission, the specializations that are brought about in vertebrate synapses, and how this can help or hinder neurological or psychiatric disorders.

## Introduction

The molecular machinery mediating neurotransmitter release is strongly conserved throughout evolution: Ca^2+^ triggers exocytosis of neurotransmitter from synaptic vesicles (SVs) in less than a millisecond by binding to Synaptotagmin, which together with Complexins activates a core fusion machinery composed of SNAREs and SV proteins (Südhof, 2013). These events are confined to specialized sites of the presynaptic plasma membrane, called active zones, by a network of proteins including RIMs, RIM binding proteins, Munc13s, CAPS and CAST/ERC proteins. One of the ways by which these scaffolding molecules act is that RIMs recruits both calcium channels and Munc13, which makes SVs tethered at the active zone fusion competent (Südhof, 2012; Imig et al., 2014; Acuna et al., 2016). The physical and functional interactions of all these proteins represent an evolutionarily conserved machinery mediating regulated neurotransmitter release at active zones. The evolutionarily conserved Ca^2+^ binding proteins Calmodulin and Synaptotagmin-7 further refine the machinery by regulating presynaptic plasticity such as short-term depression and short-term enhancement to transmitter release (Junge et al., 2004; Jackman et al., 2016).

A remarkably small number of presynaptic proteins occur exclusively in vertebrates, including the SV-associated protein Synuclein, the double C2 domain protein Doc2, the active zone scaffolding proteins Bassoon and Piccolo, and the Bassoon-binding SV protein Mover. Alpha-Synuclein appears to aid SNARE complex formation as a chaperone and its dysfunction is a hallmark of diseases called synucleinopathies, most notably Parkinson’s disease (Burré et al., 2010; Burré, 2015). Bassoon and Piccolo are very large scaffolding molecules, consisting of approximately 4000 and 5000 amino acids, respectively (Cases-Langhoff et al., 1996; tom Dieck et al., 1998; Wang et al., 1999). They do not seem to be essential for the process of transmitter release, but maintain synaptic integrity by reducing the degradation of presynaptic molecules by proteasomes and autophagy (Waites et al., 2013; Okerlund et al., 2017). As scaffolding proteins that appear very early at nascent active zones and bind a large number of proteins they may also act to recruit regulatory molecules to active zones (Fejtova and Gundelfinger, 2006; Schoch and Gundelfinger, 2006).

In a yeast-2-hybrid assay we identified the SV phospho-protein Mover – also called SVAP-30 and TPRGL (Burré et al., 2006; Antonini et al., 2008) - as a binding partner for Bassoon (Kremer et al., 2007; Ahmed et al., 2013). Mover is a small, 266 amino acid, vertebrate-specific protein. Its distribution is remarkably heterogeneous. For example, inhibitory synapses in the hippocampal CA3 region lack Mover, while excitatory synapses in the same region contain Mover (Kremer et al., 2007). Quantitative analysis revealed that the levels of Mover relative to the number of SVs varies among synapses throughout the brain (Wallrafen and Dresbach, 2018). To test how the absence of a small vertebrate-specific protein affects synapses, we investigated hippocampal synapses and the behavior of Mover knockout mice. We assumed that synapse function would not be abolished, but a modulatory role would emerge. We found that the absence of Mover affects short-term plasticity in the hippocampal CA3 but not in CA1. We show that this effect is age- and Ca^2+^-dependent, and relies on the Kainate receptor (KAR) / cyclic adenosine monophosphate (cAMP) pathway in the mossy fiber synapses. Despite increased facilitation at mossy fiber terminals, KO mice showed no impairment in spatial memory, but displayed reduced anxiety.

## Materials and Methods

### KO generation and genotyping

All animal experiments were performed in accordance with the guidelines for the welfare of experimental animals issued by the State Government of Lower Saxony, Germany. All mice (*Mus musculus*) were bred and maintained at the central animal facility of the University Medical Center, Göttingen. Embryonic stem cells (129/Ola) harboring the recombined Mover locus were generated by PolyGene (Switzerland), and injected into blastocysts. Mice with the targeted locus were crossed with a Flp deleter line to remove the Neo cassette. The resulting conditional knockout mice were crossed with mice expressing Cre recombinase under the E2A promoter to generate a global Mover knockout (KO) which were compared to its wildtype (WT) littermates. After removing Cre by breeding, the mice were back-crossed to C57BL/6J for seven generations.

Mice were routinely genotyped by PCR using genomic DNA extracted from tail biopsies using a DNA extraction kit from NextTec (Germany). For amplification the following oligonucleotide primers were used: forward primer P1 (5’-CCAATCACAAGGCGAACGAG-3’); forward primer P2 (5’-cattcagtgggacaagcaga −3’); reverse primer P3 (5’-cttggatcaggagagccttg −3). The PCR reaction was carried out for 40 cycles with denaturation at 95°C for 30 s, annealing at 56°C for 1 min, and extension at 72°C for 1 min. WT and KO animals were identified by the presence of a specific 867 bp and a 697 bp band, respectively. Initially, the KO was verified by purifying and sequencing the 697 bp band.

### Slice Preparation and Electrophysiology

Acute transverse hippocampal slices 400 µm thick were prepared from male and female 18-22 days old mice unless otherwise noted. Hippocampi were isolated and cut in a Thermo Scientific HM650V vibrotome in a cutting solution containing (in mM): 215 sucrose, 2.5 KCl, 20 glucose, 26 NaHCO3, 1.6 NaH2PO4, 1 CaCl2, 4 MgCl2, and 4 MgSO4. After sectioning, the slices were incubated for thirty minutes in a solution comprised of 50% cutting solution and 50% recording solution, which contained (in mM): 124 NaCl, 2.5 KCl, 26 NaHCO3, 1 NaH2PO4, 2.5 CaCl2, 1.3 MgSO4, and 10 glucose; 300 mOsm/kg. Recording solution for whole-cell patch clamp recordings, was comprised of (in mM): 119 NaCl, 2.5 KCl, 26 NaHCO3, 1 NaH2PO4, 4 CaCl2, 4 MgSO4, and 10 glucose; 300 mOsm/kg. After the aforementioned 30 minutes incubation, the mixed solution was changed to 100% recording solution. After changing of solution, slices were incubated for at least 60 min at room temperature. All solutions were continuously gassed with carbogen (95% O2, 5% CO2).

Excitatory post synaptic potentials (EPSPs) and receptor excitatory postsynaptic currents (EPSCs) were recorded using a HEKA EPC-10 amplifier connected to a chlorided silver wire in a borosilicate glass pipette. Recording pipettes had 0.5-1.5 MOhm pipette resistance and were filled with 1 M NaCl for extracellular recordings, and 2.0-4.0 MOhm for whole-cell recordings with a solution containing (in mM): 123 Cs-gluconate, 8 NaCl, 10 HEPES, 10 Glucose, 10 BAPTA, 5 ATP-Mg, 0.4 GTP-Na; 300 mOsm/kg, pH 7.2. Stimulation of axons was delivered through a patch-type pipette connected to either a Model A395 linear stimulus isolator (World Precision Instruments) or a Model DS3 isolated current stimulator (Digitimer).

Schaffer collateral responses were recorded at the stratum radiatum of the CA1, with the stimulation electrode placed rostral to the recording electrode. Mossy fiber field EPSPs (fEPSPs) were recorded in the stratum lucidum of the hippocampus, whereas whole-cell recordings were performed by patching CA3 pyramidal cells. In both cases the MFs were stimulated at the border between the dentate gyrus and the hilus. At the end of each MF experiment, the group II metabotropic glutamate receptor agonist DCG-IV (2S,20R,30R-2-[20,30-dicarboxycyclopropyl]glycine) was applied to the bath (1 µM) to selectively block mossy fiber responses (Kamiya et al., 1996). Recordings in which responses did not reduce by at least 80% were excluded from the analysis. For whole-cell recordings 100 µM picrotoxin was added to the solution to avoid inhibitory transmission. For each sample, every recording was repeated at least 3 times and traces were averaged. Data was sampled at 20 kHz or 50 kHz and low-pass filtered at 2.9 kHz.

### Behavior

The cross-maze, Morris water maze, open-field and elevated plus maze tests were performed as previously described (Jawhar et al., 2012; Bouter et al., 2013) and are further described in detail. Only male mice were used for behavior experiments. The cross-maze built from black plastic consists of four arms (30 cm length × 8 cm width × 15 cm height) arranged in 90° position extending from a central region (8 cm length × 8 cm width × 15 cm height). During a 10 minute test session, each mouse was randomly placed in one arm and allowed to freely explore the maze. Alternation was defined as successive entries into the four arms in overlapping quadruple sets. The alternation percentage was calculated as the percentage of actual alternations to the possible number of arm entries. To diminish odor cues, the maze was cleaned with 70% ethanol solution between mice.

The elevated plus maze consists of two open and two closed arms (15 cm length × 5 cm width) extending at 90° angles from a central area (5 cm length × 5 cm width) raised 75 cm above a padded surface. During the test the mouse is placed onto the center field facing one of the open arms and is allowed to explore the maze for 5 min. Anxiety was measured by the time spent in the open arms, with lower anxiety levels corresponding to more time spent in open arms.

In the open-field the mice were tested for 5 min in a 50 × 50 cm arena with 38 cm-high walls. Time spend in the center of the maze correlates to lower anxiety levels. Every maze described above was cleaned with 70 % ethanol different trials to diminish odor cues between mice.

In the Morris water maze, mice had to learn to use spatial cues to locate a hidden, circular platform (10 cm) in a circular pool (110 cm diameter) filled with tap water made opaque by non-toxic white paint. Water was maintained at 20 °C for the test duration. Testing began with 3 days of cued training. For these trials, the platform was marked with a triangular flag. Mice were introduced into the water at the edge of the pool facing the wall. They were then given 1 min to find the submerged platform. Mice that failed to find the platform in 60 s were gently guided to it. All mice were allowed to sit on the platform for 10 s before being removed from the pool. To prevent hypothermia, all mice were kept in front of a heat lamp until the next trial. Each mouse received four training trials per day with an average inter-trial interval of 15 min. Both the location of the platform and the position at which mice were introduced into the pool changed between trials.

Twenty-four hours after the last day of cued training, mice performed 5 days of acquisition training, in which the flag was removed from the platform. In addition to the distal cues existing in the room, proximal visual cues were attached to the outside of the pool. The platform location remained stationary for each mouse throughout training. At the start of every trial, mice were introduced into the pool from one of four predefined entry points. The order in which these entry points were used varied between training days. Trials were conducted as during the cued training phase. Twenty-four hours after the last acquisition trial, a probe test was performed to assess spatial reference memory: the platform was removed from the pool and mice were introduced into the water from a novel entry point. Mice were then allowed to swim freely for 1 min while their swimming path was recorded.

### Experimental Design and Statistical Analysis

Electrophysiological data was analyzed using custom-written procedures in Igor Pro 6.3 (Wavemetrics). ANY-Maze video tracking software (Stoelting Co.) was used to record the parameters of behavioral experiments. Statistical significance was tested using GraphPad Prism 7.04 (GraphPad Software). 2-way ANOVA tests were used to analyze high-frequency trains of stimulation and paired-pulse ratios; extra sum-of-squares F test was used to test differences between curve fits. For all other tests two-tailed Student’s unpaired T test was used. For every experiment 3 or more animals were used. Results are reported as mean ± SEM whereas “n” refers to the number of slices recorded and “N” the number of animals.

## Results

To obtain a global knockout of Mover we bred Mover conditional knockout mice with E2A-Cre mice (Fig. 1A). The E2A promotor drives Cre expression in the early mouse embryo, thus excising Mover in all cells from early embryonal stages on. The entire Mover gene consists of less than 4000 base pairs, including 4 exons and 3 introns. We verified the expected excision of Mover exons 1, 2 and 3 by PCR (Fig. 1B), and by sequencing the PCR product (Fig. 1C). Western blotting revealed that Mover was not detected in brain homogenates from Mover knockout mice (Fig. 1D). Likewise, there was no Mover immunofluorescence in sections of the hippocampus from Mover knockout mice (Fig. 1E).

**Fig. 1:**
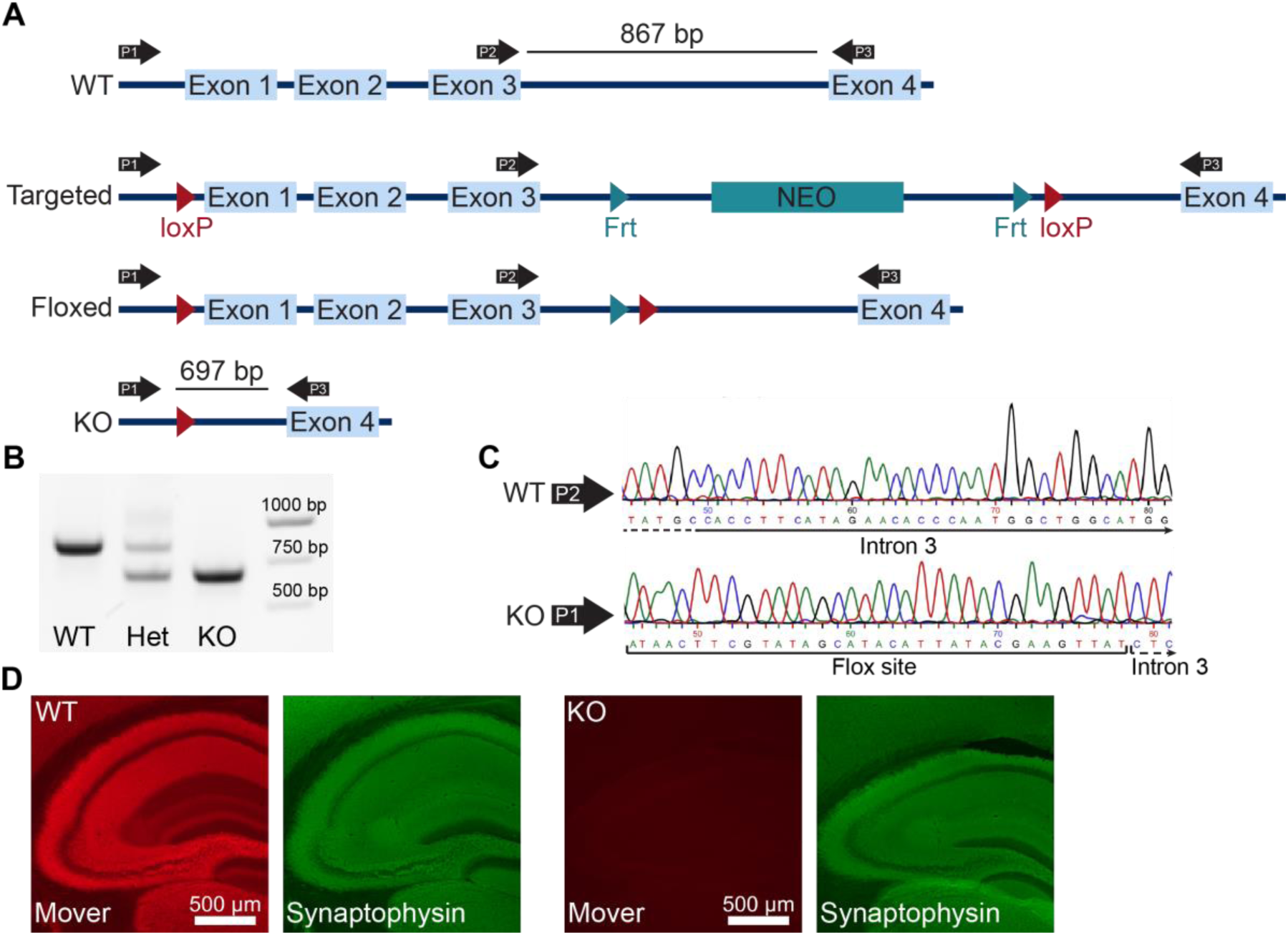
Global knockout of Mover. (A) Gene targeting strategy for Mover KO mice. (B) Results of the PCR used for genotyping. Primers P1, P2 and P3 shown in (A) were always used in the same reaction. When a WT and a KO allele were present, P1 and P3 produce a 697 bp band, P2 and P3 produce a 867 bp band (lane “Het”). When only WT alleles are present the primers produce only the 867 bp band (lane “WT”), when only KO alleles are present the primers produce only the 697 (lane “KO”). (C) Example of sequencing results for WT (top) and KO (bottom). Examples shown start from nucleotide 45 from sequencing result and show a part of intron 3 using the primer P2 for WT and the flox site followed by Intron 3 in KO, showing the absence of exons 1 to 3. (D) Immunofluorescence of WT (left) and KO (right) mouse brain sections stained for Mover and Synaptophysin.

### Mover knockout affects mossy fiber but not Schaffer collateral synaptic transmission

The absence of Mover in inhibitory synapses of the hippocampus (Wallrafen and Dresbach, 2018), together with its layered structure, makes it an ideal target to study the effect of Mover knockout in glutamatergic transmission. Recording of field excitatory post synaptic potentials (fEPSP) in the hippocampal CA1 with stimulation of the Schaffer collaterals (SC) with increasing stimulation strength allows for an assessment of synaptic strength based on input-output curves. Input-output curves showed no difference in synaptic strength between WT and KO mice (Fig. 2A_1_).

**Fig. 2:**
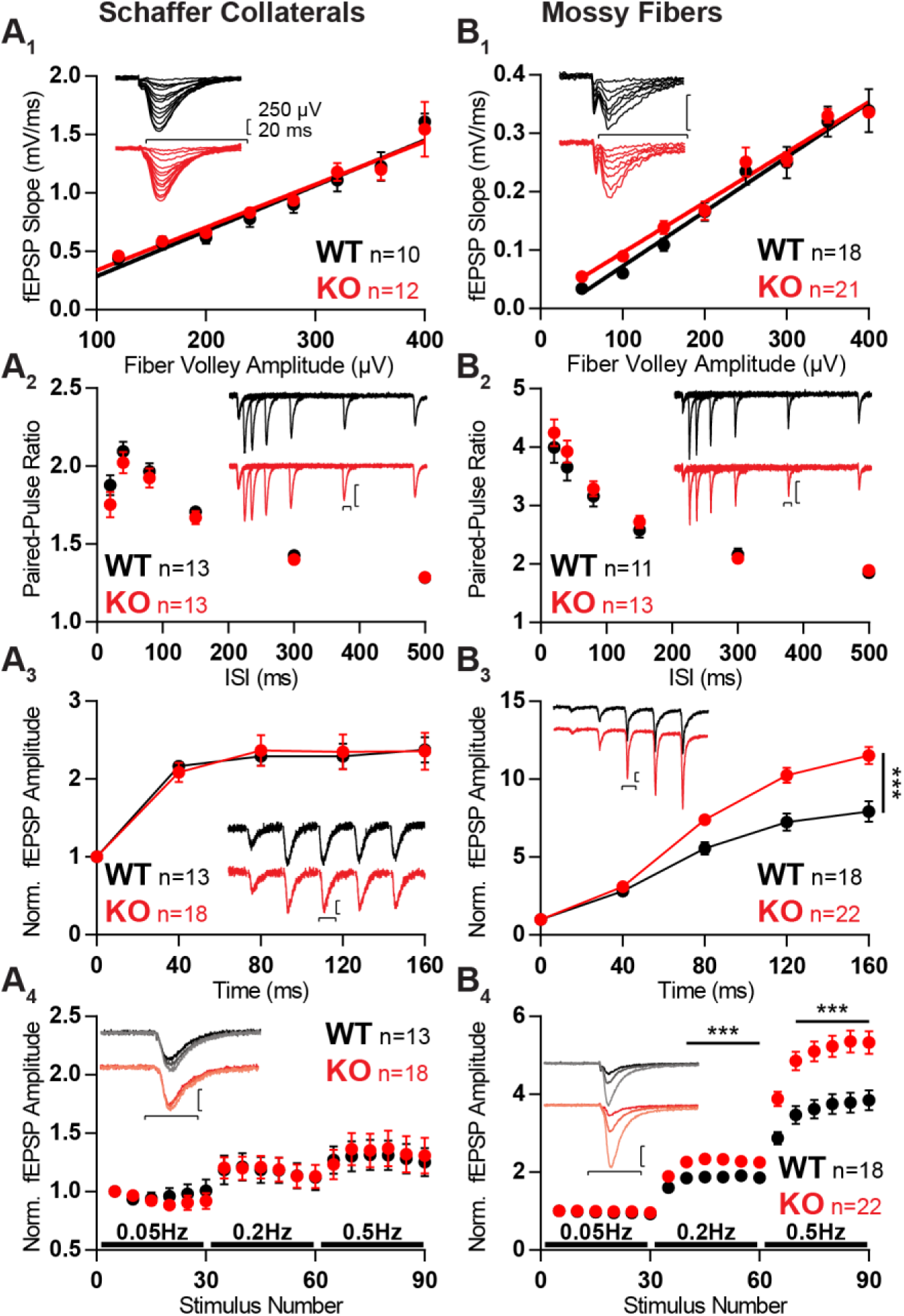
Synaptic transmission unchanged in SC in absence of Mover (*A*), but increased activity-dependent transmission in MF (*B*). (*A_1_*, *B_1_*) fEPSP slopes versus fiber volley recorded at increasing stimulation strengths depict an input-output relationship unchanged by the absence of Mover. (*A_2_*, *B_2_*) Paired-pulse ratios recorded from fEPSP amplitudes across varying inter-stimulus intervals show no difference between WT and KO. (*A_3_*, *B_3_*) Normalized responses to a 25 Hz train of stimulation showed increased facilitation in KO MF when compared to WT, but not in SC. (*A_4_*, *B_4_*) Normalized responses to stimuli delivered at 0.05, 0.2 and 0.5 Hz reveal increased facilitation in KO MF. Each dot represents the average response to 5 consecutive stimuli. Responses were normalized to the amplitude of the first fEPSP. (*Insets*) Representative traces from WT (*black*) and KO (*red*) hippocampi. Scale bars: vertical=250 µV, horizontal=20 ms. Error bars indicate SEM. SC WT n= 10-13; SC KO n=12-18; MF WT n=11-18; KO n=13-22 ***: *p*<0.001.

To further probe changes in presynaptic transmission we evaluated short-term plasticity with three different protocols: paired-pulse facilitation with two pulses given at different inter-stimulus intervals (ISI; Fig. 2A_2_), burst-induced facilitation with a 25 Hz train consisting of 5 stimuli (Fig. 2A_3_), and low-frequency facilitation by increasing frequency of stimulation from 0.05 Hz to 0.2 Hz and to 0.5 Hz (Fig. 2A_4_). In all protocols no differences between WT and KO were detected.

**Fig. 3:**
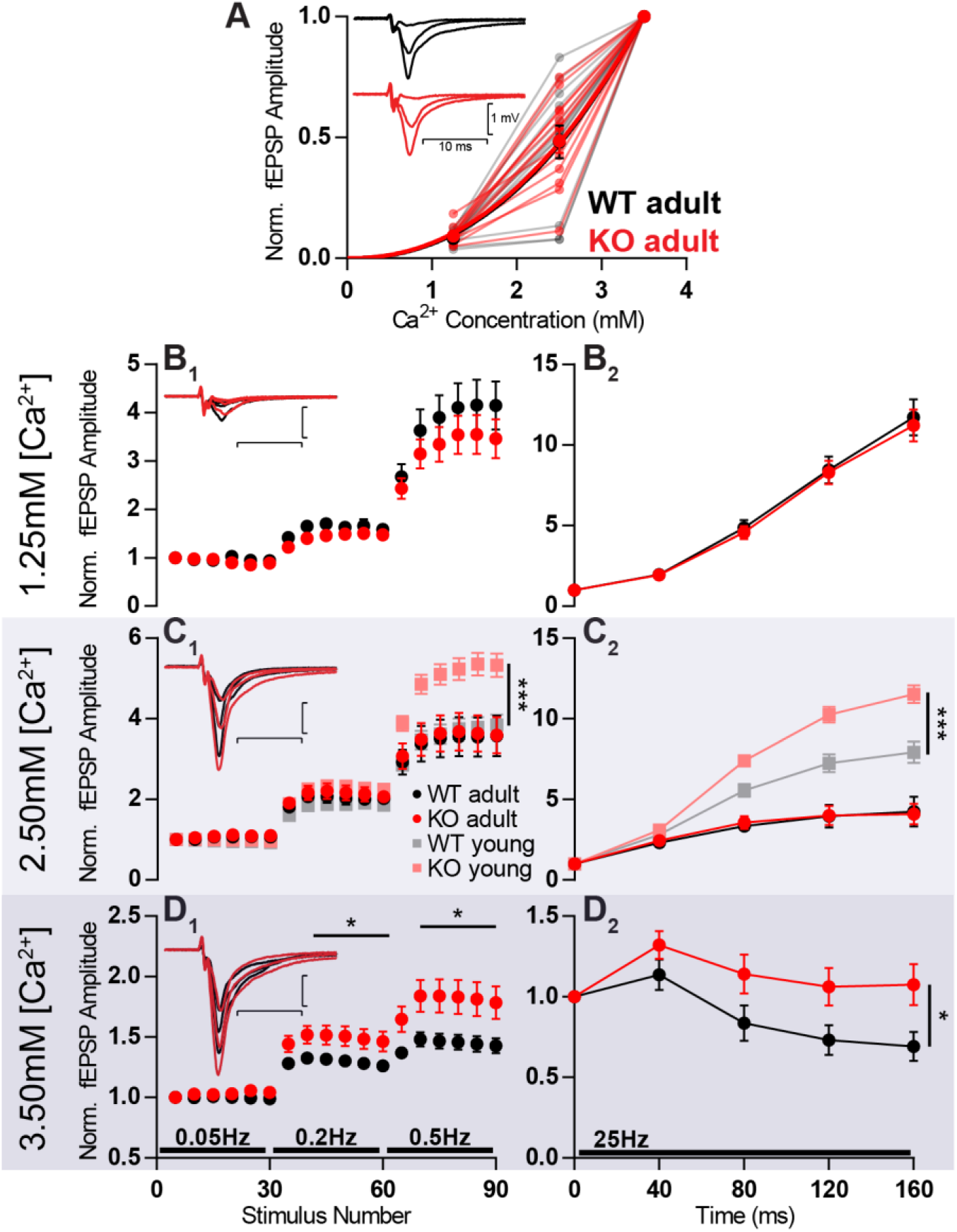
Increased facilitation in KO is age- and calcium-dependent. In MF of 8-week old animals KO (KO adult) has stronger facilitation than WT (WT adult) only in high extracellular calcium concentration. (*A*) Increasing calcium concentration leads to similar baseline responses from WT and KO. Responses were normalized to fEPSC amplitudes at 3.5 mM Ca^2+^. (*B-D*) Short-term plasticity at different extracellular calcium concentrations. (*B_1_*) Normalized responses to stimuli delivered at 0.05, 0.2 and 0.5 Hz at 1.25 mM extracellular calcium. (*B_2_*) Normalized responses to a 25 Hz train of stimulation at 1.25 mM extracellular calcium. (*C_1_*) Normalized responses to stimuli delivered at 0.05, 0.2 and 0.5 Hz at 2.5 mM extracellular calcium, in four different conditions: 8-week old WT (adult), 8-week old KO (adult), 3-week old WT (WT young), 3-week old KO (KO young). The last two conditions are the same dataset as present in Fig. 1B_3_-B_4_. (*C_2_*) Normalized responses to a 25 Hz train of stimulation at 2.5 mM extracellular calcium. (*D_1_*) Normalized responses to stimuli delivered at 0.05, 0.2 and 0.5 Hz at 3.5 mM extracellular calcium. (*D_2_*) Normalized responses to a 25 Hz train of stimulation at 3.5 mM extracellular calcium. (*B_1_, C_1_, D_1_*) Each dot represents the average response to 5 consecutive stimuli. (*B-D*) Responses were normalized to the amplitude of the first fEPSP. (*Insets*) Representative traces from WT (*black*) and KO (*red*) hippocampi. Scale bars: vertical=1 mV, horizontal=10 ms. Error bars indicate SEM. WT adult n=13; KO adult n=13. *: *p*<0.05; ***: *p*<0.001.

**Fig. 4:**
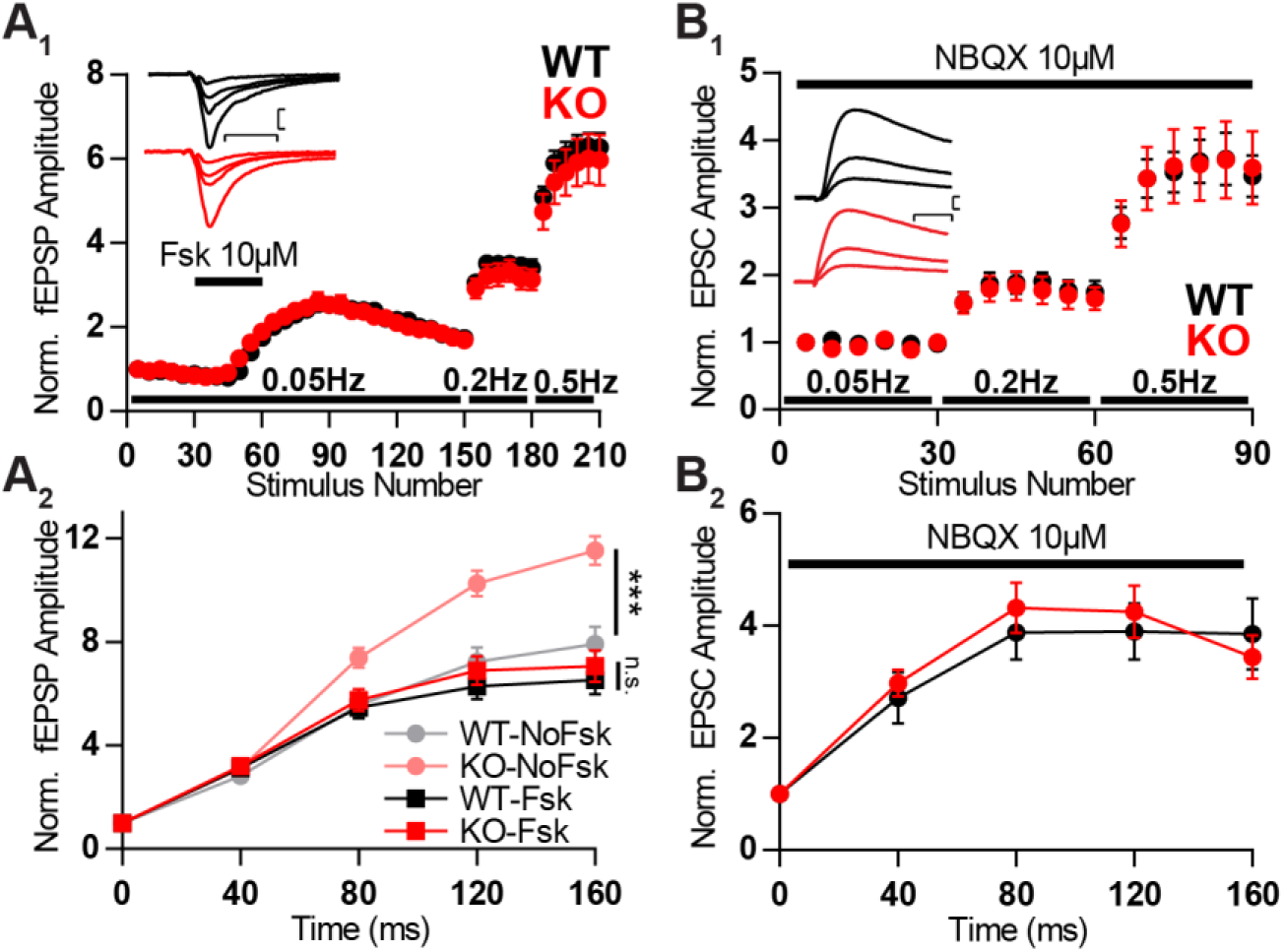
Forskolin potentiation and receptor blockade by NBQX occlude KO boost in facilitation. (*A_1_*) Normalized MF fEPSP amplitudes during the time course of experiment in which forskolin (*fsk*, 10µM) is applied for 10 minutes and frequency of stimulation is changed from 0.05 to 0.2 and further to 0.5 Hz. (*A_2_*) Normalized MF responses to a 25 Hz train of stimulation in four different conditions: WT without forskolin application (WT-NoFsk), KO without forskolin application (KO-NoFsk), WT after forskolin potentiation (WT-Fsk), and KO after forskolin potentiation (KO-Fsk). The first two conditions are the same dataset as present in Fig. 1 B_3_. (*B*) NMDA EPSCs from MF stimulation and recording from CA3 pyramidal cells in the presence of 10 µM NBQX. (*B_1_*) Normalized responses to stimuli delivered at 0.05, 0.2 and 0.5 Hz. (*B_2_*) Normalized responses to a 25 Hz train of stimulation in the presence of 10µM NBQX. (*A_1_*,*B_1_*) Each data point corresponds to the average response to 5 consecutive stimuli. (*Insets*) Representative traces from WT (*black*) and KO (*red*) hippocampi. Responses were normalized to the amplitude of the first response. (*A*) Scale bars: vertical=200 µV, horizontal=10 ms; WT n=10; KO n=10. (*B*) Scale bars vertical=50 pA, horizontal=10 ms; WT n=9 KO n=10. Error bars indicate SEM. *n.s.*: not significant; ***: *p*<0.001.

With no apparent differences in SC, we moved to investigate the CA3, more specifically the CA3 pyramidal cell inputs from mossy fiber terminals (MF). Similarly to SC, input-output curves and paired pulse facilitation were unchanged by the absence of Mover (Fig. 2B_1,2_). However, the use of repeated stimulation, either by bursts (*p*=0.0004) or by low-frequency stimulation (0.05Hz vs 0.2Hz: *p*=0.0007; 0.05Hz vs 0.5Hz: *p*=9.9e-05), leads to a much stronger facilitation in KO than in WT (Fig. 2B_3,4_). KO MF facilitated almost 50% more than WT. Therefore, Mover, when present, restricts short-term plasticity in the CA3 but not in CA1.

### Mover affects MF facilitation under high calcium conditions

The extent of facilitation in MF is known to vary with age (Mori-Kawakami et al., 2003). Hence, to further explore the effect of the absence of Mover we analyzed the effect of MF stimulation in older mice (8-week old) with different extracellular calcium concentrations. The change in calcium concentration was compensated with magnesium, as to keep the concentration of divalent ions constant.

Firstly, increasing the concentration of calcium from 1.25 mM to 2.5 mM and further to 3.5 mM led to an increase in fEPSP amplitude. This increase was similar when comparing WT and KO responses (Fig. 3A).

When testing the previously used short-term plasticity protocols, at 1.25 mM Ca^2+^ the extent of facilitation did not differ between WT and KO anymore (Fig. 3B). Surprisingly, after increasing Ca^2+^ to 2.5 mM, WT and KO continued to facilitate to the same extent, whereas the same calcium concentration in 3 week-old animals promoted a stronger facilitation in MF KOs (Fig. 3C, Fig. 2B_3,4_). This reveals an age-dependency of the effect that Mover has on facilitation.

When further increasing Ca^2+^ concentration to 3.5 mM, the difference in the extent of facilitation between KO and WT becomes obvious again for both high-frequency (*p*=0.04; Fig. 3D_2_) and low-frequency facilitation (0.05Hz vs 0.2Hz: *p*=0.04; 0.05Hz vs 0.5Hz: *p*=0.03; Fig. 3D_1_). The KO responses facilitate more than WT, corroborating what was observed in younger animals and the idea that Mover acts in a calcium-dependent manner.

### Forskolin and NBQX occlude boost in facilitation observed in KO

Synaptic plasticity in MF is known to be strongly tied to intracellular levels of cAMP and is, therefore, subject to regulation by forskolin (Weisskopf et al., 1994). Hence, since both forskolin and Mover are strongly affecting plasticity in MF we decided to investigate if Mover acts in the cAMP pathway.

We anticipated that, if Mover participates in this pathway and is involved in this long-term potentiation, we would observe changes in the degree of potentiation caused by forskolin. If on the other hand, Mover is not involved in the expression of the potentiation, but would inhibit the pathway upstream of cAMP, no differences in Forskolin-potentiation would be observed. That was indeed the case: application of forskolin while recording MF responses led to a similar potentiation in WT and KO fEPSCs (Fig. 6A_1_). However, if Mover is inhibiting the pathway prior to the synthesis of cAMP, application of forskolin would occlude any effect of the presence or absence of Mover. Correspondingly, after forskolin potentiation we observed a lack of difference between WT and KO low- and high-frequency facilitation (Fig. 4A_1,2_), which differed from what we have previously shown (Fig. 2B_3,4_, also in Fig. 4A_2_ for comparison). Therefore, potentiation by forskolin occluded the increased facilitation observed in the KO.

**Fig. 5.**
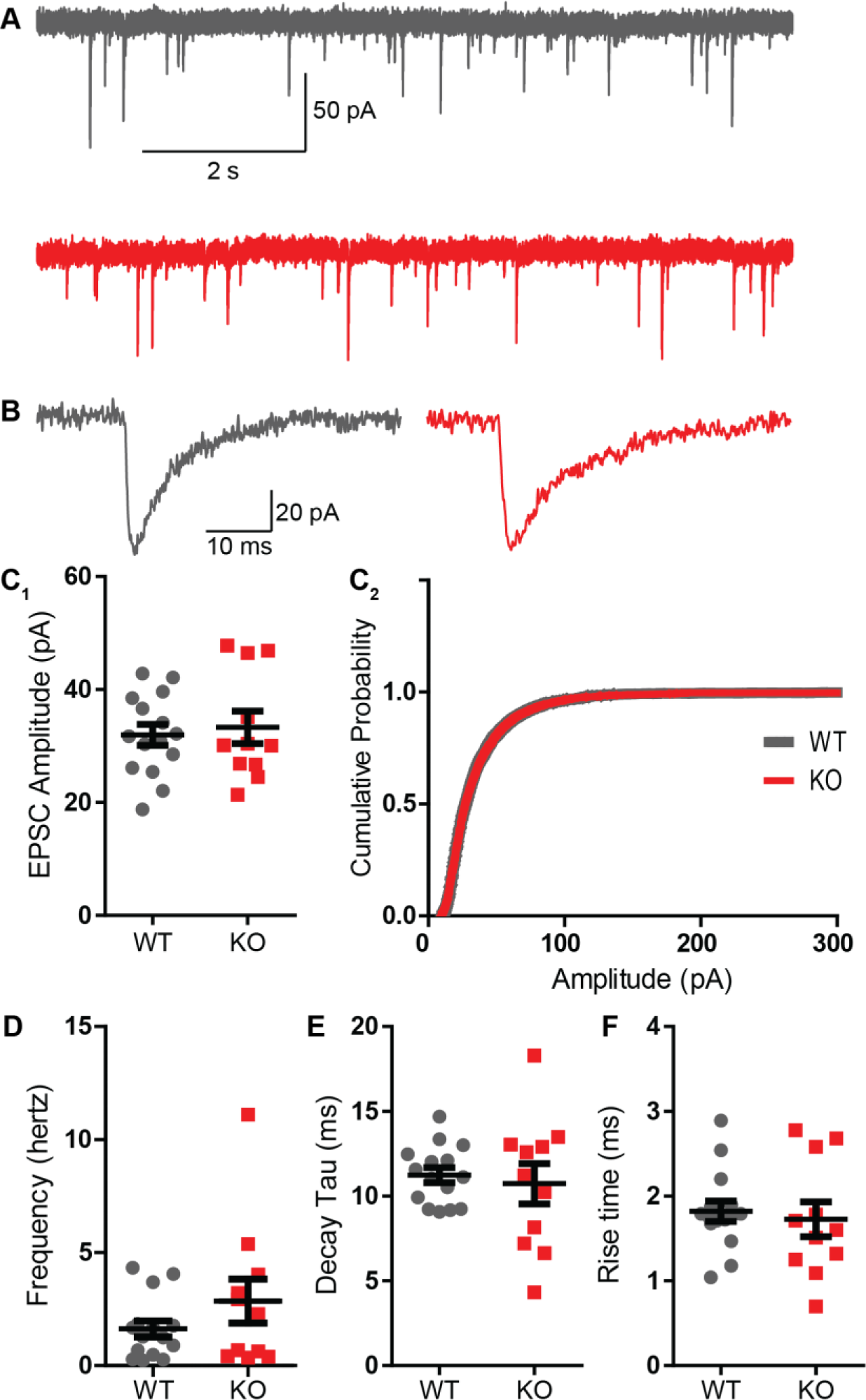
Absence of Mover does not interfere with miniature EPSC parameters in CA3 pyramidal cells. (*A*) Representative traces from WT (grey) and KO (red) CA3 pyramidal cells under presence of 1µM TTX. (*B*) Representative miniature EPSC waveform from traces in A. (*C*) Amplitudes of miniature EPSC events were unchanged in their average amplitude (*C1*) and in their cumulative probability (*C2*). (*D*) Frequency of events was not changed by the absence of Mover. Miniature EPSC dynamics, namely the time constant of decay (*E*) and the 10-90 rise time (*F*), showed no difference between WT and KO. Error bars indicate SEM. WT n=15; KO n=11.

**Fig. 6:**
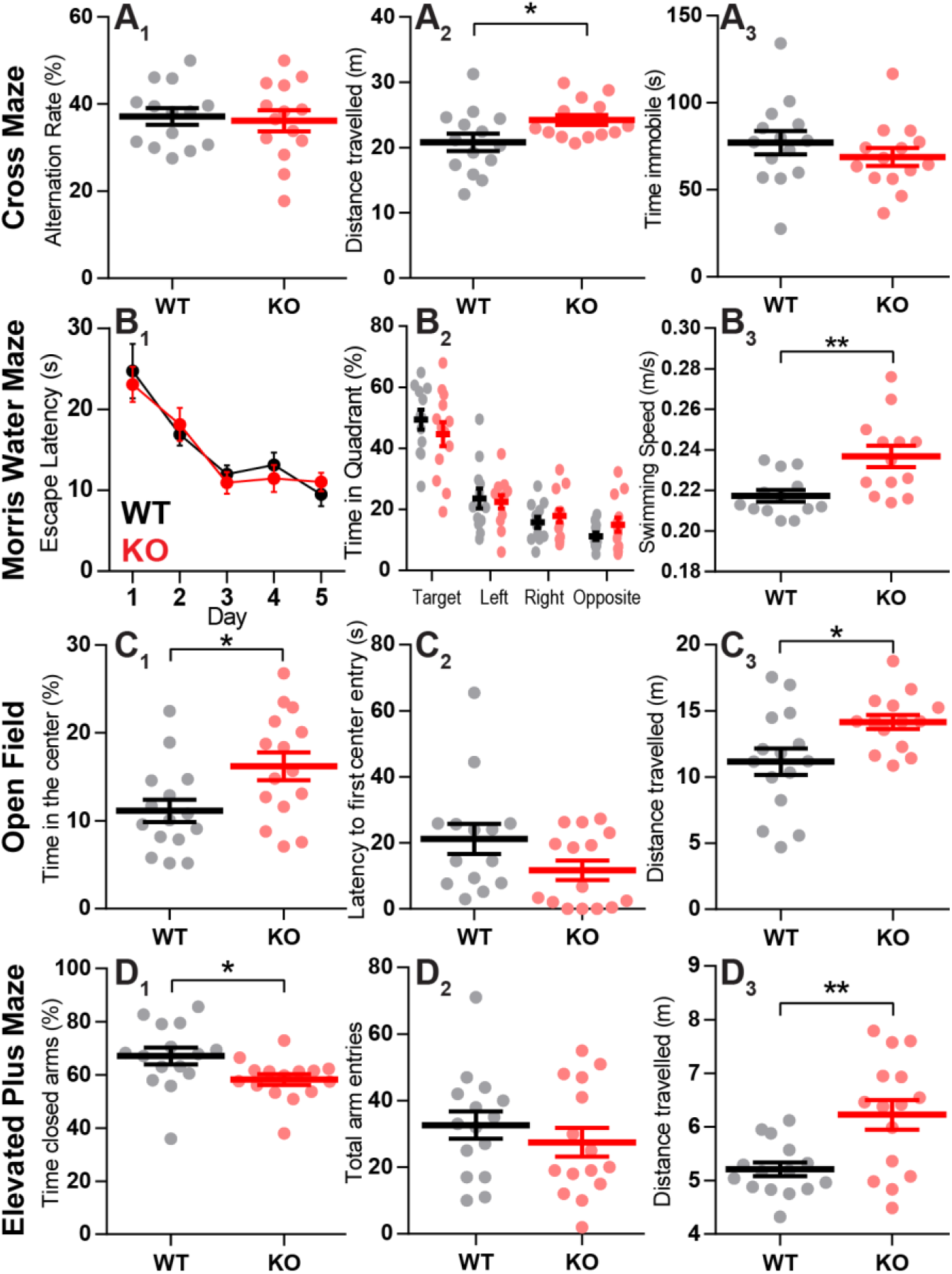
KO mice display unimpaired spatial reference memory but increased exploratory behavior. (*A*) Mice were tested on their working memory based on the alternation rate of arm entries in a cross-maze. Alternation rate (*A_1_*) and immobility time (*A_3_*) were not different between WT and KO, but KO travelled longer distances (*A_2_*). (*B*) Long-term spatial reference memory was assessed with the Morris water maze. (*B_1_*) Across 5 days of acquisition animals of both genotypes showed a similar decrease in escape latencies to reach the hidden platform. (*B_2_*) During probe trial, where the platform was removed, time spent in each quadrant did not vary between WT and KO. However, both genotypes spent significantly more time in the target quadrant. (*B_3_*) During probe trial KO mice showed increased swimming speed over WT. (*C*) Anxiolytic effect observed in KO mice in open-field test. (*C_1_*) KO mice spend significantly more time in the center area of the open-field arena than WT. (*C_2_*) WT and KO mice display similar latencies to their first visit to the center area. (C_3_) KO mice travelled longer distances than WT. (*D*) KO animals display reduced anxiety-like behavior in the elevated plus maze. (*D_1_*) KO mice spend less time in closed arms than WT. (*D_2_*) WT and KO mice had a similar number of arm entries in general. (*D_3_*) KO mice walked longer distances than their WT counterpart. Error bars indicate SEM. WT N=14-15; KO N=12-15.*: *p*<0.05; **: *p*<0.01.

The activation of KARs in MF have also been reported to participate in presynaptic plasticity in the cAMP pathway (Bortolotto et al., 1999; Kamiya et al., 2002; Lauri et al., 2003; Rodríguez-Moreno and Sihra, 2004; Scott et al., 2008; Andrade-Talavera et al., 2012). To further evaluate this possibility we used whole-cell intracellular recordings of CA3 pyramidal cells. Basic synaptic transmission measured by whole cell recordings was unchanged by the absence of Mover: miniature EPSCs, recorded with the presence of 1 µM tetrodotoxin (TTX), had similar frequency, amplitude and kinetics between WT and KO (Fig. 5).

To evaluate KAR participation we blocked KAR and α-amino-3-5-methyl-4-isoxazolepropionic acid (AMPA) receptors with the use of 10µM 2,3-dihydroxy-6-nitro-7-sulfamoylbenzo[f]quinoxaline-2,3-dione (NBQX). Responses comprised of N-methyl-D-aspartate (NMDA) EPSCs. Similarly to the experiments with forskolin, the use of NBQX occluded any effect of the KO in short-term plasticity: WT and KO facilitated to the same extent both in low-frequency facilitation (Fig. 4B_1_) and in high-frequency trains (Fig. 4B_2_). Taken together, these results suggest that Mover plays a role in the pathway of KAR and cAMP in MF, buffering facilitation in situations of high activity.

### Absence of Mover does not affect memory but has anxiolytic effect

The hippocampus is strongly involved in learning and memory via its trisynaptic circuit. But would an increase in short-term plasticity in the MF terminals affect memory? The influence on synaptic plasticity in CA3 but not on CA1 by the absence of Mover led to the question whether memory in KO mice was affected. Short-term plasticity, as well as MF synaptic transmission, have been implicated in memory and cognition (Gilbert and Kesner, 2006; Kesner, 2007; Liu et al., 2013). We therefore, explored the influence of the absence of Mover on memory with two different experiments: the cross-maze and Morris water maze.

Working memory was assessed by analyzing the mice spontaneous alternation between arms of the cross-maze. Working memory is unaltered in the absence of Mover, as alternation rates did not vary between WT and KO (Fig. 6A_1_). Mice also had similar amount of immobility time, however KO mice travelled longer distances than WT (*p*=0.036; Fig. 6A_2,3_).

Spatial reference memory of mice was evaluated using the Morris water maze. Firstly, mice underwent a cued training to familiarize with the pool, ensure that they had intact vision and the appropriate motor abilities to swim. Secondly, across the 5 days of acquisition training animals of both genotypes showed a similar decrease in the escape latency to reach the hidden platform (Fig. 6B_1_).

Twenty-four hours after the last acquisition trial, a probe trial was performed to assess spatial reference memory. WT and KO mice displayed similar preference for the target quadrant, as indicated by the percentage time spent in different quadrants of the pool (Fig. 6B_2_). This indicates that spatial reference memory is unaffected by the absence of Mover. Furthermore, during the probe trial KO mice swam significantly faster than WT mice (*p*=0.006; Fig. 6B_3_).

The observed increase in exploratory behavior observed by increased distance travelled in the cross-maze and increase in swimming speed in the water maze indicate a possible anxiolytic effect in the absence of Mover. To test anxiety levels in the mice we made use of two different experiments: the open-field test and the elevated plus maze.

Exploratory and spontaneous locomotor activity of KO mice was compared to WT mice in the open-field test. KO mice spent significantly more time in the center of the maze in comparison with their WT littermates (*p*= 0.019; Fig. 6C_1_) reflecting reduced anxiety. There was no significant difference in the time mice took to reach the center of the arena at the beginning of the experiment. Furthermore, confirming results from the previous experiments, KO mice travelled significantly more than WT (*p*= 0.014; Fig. 6C_3_). The results in the elevated plus maze further corroborated the findings from the open-field test. KO mice showed a decreased anxiety phenotype which was indicated by spending less time in the closed arms (*p*=0.028; Fig. 6D_1_). Furthermore, KO mice travelled significantly more than WT mice (*p*=0.002; Fig. 6D_3_), while total number of arm entries was unchanged (Fig. 6D_2_).

## Discussion

Mover is a vertebrate-specific protein associated with synaptic vesicles. The fact that invertebrates do not require Mover for synaptic transmission suggests that Mover has a modulatory role, and may regulate the conserved molecular machinery underlying neurotransmitter release.

To test whether Mover has a role in transmitter release we generated a mouse line lacking Mover. We found that the knockout of Mover affects short-term synaptic plasticity in the hippocampal MF but not in the downstream synapses, i.e. SC. In particular, frequency facilitation, a hallmark of presynaptic plasticity at MF (Nicoll and Schmitz, 2005), was increased in Mover knockout mice. High-frequency trains of stimulation also led to stronger facilitation in the absence of Mover. This increased facilitation was stronger in younger animals and in situations of high calcium concentration, and it was occluded by increasing cAMP levels or by blockade of KARs.

Remarkably, the absence of Mover did not lead to a decline in performance during tasks that rely on spatial memory. However, KO mice showed increased locomotor activity, which often correlates with lower anxiety levels (Liebsch et al., 1998). Indeed, results with anxiety-related tasks show anxiogenesis correlated with Mover presence.

### The role of Mover in synaptic release is dependent on the calcium concentration in the presynaptic terminal

In principle, the increase in synaptic facilitation observed at MF could arise from a reduction of the initial release probability. In the current study, however, we see no indication of a change in basal release, as indicated by the lack of change in paired-pulse ratio, input-output, basal responses to increasing calcium concentration and EPSC amplitude. On the other hand, our results suggest that in the KO there is an activity-dependent increase in neurotransmitter release, likely due to calcium influx. Two observations suggest that the increase in facilitation observed in the Mover KO arises in conditions of high intracellular calcium. First, under conditions of high extracellular calcium, KO MF showed increased facilitation compared to WT. That was not the case when extracellular calcium concentrations were lower. Second, the effect on facilitation only occurred during repetitive activity, which leads to an intraterminal buildup of calcium.

The idea that Mover KO leads to an increase of neurotransmitter release in MF corroborates results in the calyx of Held, where knockdown of Mover led to a reduction in the intrinsic calcium-sensitivity of release (Körber et al., 2015). Taken together these results suggest that Mover constrains release, under conditions of high Ca^2+^ in MF and under basal conditions in the calyx of Held. Both the change in calcium-sensitivity observed at the Calyx of Held and the calcium-dependency of Mover action at MF suggest that Mover is part of a calcium-sensing pathway.

### Linking Mover, Calcium, cAMP and Kainate receptors

How does Mover interact with calcium signaling? The amino acid structure of Mover does not reveal any canonical calcium binding domain (Kremer et al., 2007). However, it is known that Mover binds to CaM in a calcium-dependent manner (Körber et al., 2015). In hippocampal MF, the rise in calcium concentration stimulates adenylyl cyclase through the binding of CaM (Rodríguez-Moreno and Sihra, 2004; Andrade-Talavera et al., 2012). There are two CaM-binding isoforms of adenylyl cyclase present in MF which are known to affect synaptic plasticity: adenylyl cyclase 1 (Villacres et al., 1998) and 8 (Wang et al., 2003). Our results suggest that Mover and adenylyl cyclase act in the same pathway: activation of adenylyl cyclase by forskolin led to occlusion of the Mover knockout effect. Mover could, therefore, act on CaM to inhibit cAMP synthesis. Furthermore, this effect would be calcium-dependent, which agrees with Mover changing short-term plasticity in high calcium and during repetitive stimulation.

During repetitive stimulation Kainate autoreceptors in MF terminals facilitate Ca^2+^ influx (Kamiya et al., 2002; Lauri et al., 2003; Scott et al., 2008; Andrade-Talavera et al., 2012). The rise in intracellular Ca^2+^ is linked to the aforementioned increase in cAMP by Ca^2+^-Calmodulin activation of adenylyl cyclase (Rodríguez-Moreno and Sihra, 2004; Andrade-Talavera et al., 2012). KARs are activated by glutamate release from the MF terminals in which they are present, thereby creating a positive feedback loop which increases synaptic transmission in this synapse in short- and long-term ranges (Lauri et al., 2001b; Andrade-Talavera et al., 2012; Negrete-Díaz et al., 2018). In this way, these receptors participate in short- and long-term plasticity in MF terminals (Lauri et al., 2001a, 2001b; Schmitz et al., 2001; Ji and Stäubli, 2002; Contractor et al., 2003). The notion that Mover constrains facilitation by inhibiting cAMP synthesis is consistent with the participation of KARs in MF plasticity: if Mover indeed inhibits cAMP synthesis, it would act downstream of KARs. Therefore, the blockade of KARs would lead to similar facilitation in WT and KO, thus ignoring the presence or absence of Mover. Consistent with this prediction, we found that the blockade of KARs by NBQX abolished the increased facilitation observed in the KOs.

The different participation of Kainate autoreceptors in SC versus MF is consistent with the different effects of the KO in these two synapses. Activation of KARs in the mossy fibers has been shown to have a wide range of effects from facilitating synaptic responses under small concentrations of kainate (Lauri et al., 2001a; Schmitz et al., 2001; Ji and Stäubli, 2002) to depressing synaptic responses when applying higher kainate concentrations (Kamiya and Ozawa, 2000; Schmitz et al., 2000). On the other hand, in SC, activation of KAR only leads to synaptic depression (Chittajallu et al., 1996; Frerking et al., 2001; Negrete-Díaz et al., 2018). Since KARs do not trigger facilitation in SC, the absence of Mover does not affect it in this synapse. Thus, our data are consistent with the idea that Mover, as a vertebrate-specific protein, and in a synapse-specific way, regulates the conserved machinery of CaM and cAMP.

### Mover could buffer synaptic strength

MF transmission relies heavily on facilitation for efficient information transfer and is, therefore, considered a ‘conditional detonator’(Vyleta et al., 2016). We have shown that Mover constrains facilitation in this synapse, possibly keeping detonation within physiological range. On the other hand, Mover reduces the amplitude of the first response to a train of stimuli and subsequent depression in a synapse where this initial response is vital for transmission of fast and reliable auditory information, the calyx of Held (Englitz et al., 2009; Körber et al., 2015). Thus, in general, Mover may act to buffer synaptic strength.

Homeostatic plasticity is a neuronal compensatory adjustment in response to overall network activity in order to keep neuronal firing rates in a physiological range (Lazarevic et al., 2013). In a study in the endbulb of Held, disruption of Bassoon led to downregulation of Mover and an increase in release probability, which was attributed to homeostatic plasticity (Mendoza Schulz et al., 2014). Since the presence of Mover seems to correlate with a reduction in release – during the first response at auditory brainstem synapses and during trains at MF - it is possible that Mover is involved in bringing about this homeostatic change. This is consistent with the idea that Mover could act to keep neurotransmitter release within healthy levels and should be investigated in the future.

### Importance of Mover for proper anxiety-responses

Alterations in synaptic plasticity in the hippocampus often lead to impairment in spatial memory (Bannerman et al., 2014). The similar time regime of MF frequency facilitation and working memory has led to the suggestion that such facilitation would be the biological substrate of working memory (Kesner, 2007; Hagena and Manahan-Vaughan, 2010). Our study argues against that assumption as the altered facilitation in KO did not lead to any change in spatial working memory. Long-term spatial memory performance in the Morris water maze was also not affected. In contrast, KO mice displayed increased exploratory behavior, which is often a sign of decreased anxiety. Indeed, the open-field test and the elevated plus maze confirmed an anxiolytic effect upon the absence of Mover. While we do not defend a causal link between the observed anxiolytic effect and the increased synaptic facilitation at MF, such a link is not an unrealistic scenario, as many studies have been surfacing linking hippocampal synaptic plasticity to anxiety (Bannerman et al., 2014). Independent of the site of action, Mover is, therefore, important to allow for proper anxiety responses. The link between the expression of Mover and anxiety is not the first to connect this protein to a psychiatric disorder: Mover has been shown to be strongly upregulated in the brains of schizophrenic patients (Clark et al., 2006). Interestingly, other synaptic proteins, such as SNARE proteins, have been implicated in schizophrenia and the respective mouse models show an anxiety phenotype (Katrancha and Koleske, 2015). Furthermore, it has been proposed that runaway excitation, possibly due to glutamate spillover, is a prominent feature of many psychiatric disorders such as schizophrenia (O’Donovan et al., 2017). This highlights the appeal of future studies addressing the role of Mover not only in synaptic transmission but also in pathophysiology, as Mover might have evolved to buffer synaptic strength and avoid runaway neurotransmitter release.

## Acknowledgements

This study was funded by the DFG Center for Nanoscale Microscopy and Molecular Physiology of the Brain (CNMPB, B1-1, to TD and CNMPB, C1, to TAB and YB), as well as by the Excellence fellowship (DFG Grant GSC 226/1) provided by the Göttingen Graduate School of Neurosciences, Biophysics and Molecular Biosciences (GGNB, to JSV) and by the Jacob-Henle-Program for Experimental Medicine of the University Medicine Göttingen (to JMW). We would like to thank Irmgard Weiß and Petra Tucholla for expert technical assistance. We would also like to thank Thomas J. Younts and Yoav Ben-Simon for help with the electrophysiological recordings as well as Tobias Moser for kindly providing Cs-gluconate and many fruitful discussions. Additionally, we thank Asha Kiran Akula, Nils Brose and Kerstin Reim for support during the generation of the conditional knockout mice. The authors declare no competing financial interests.

Author contributions
**J.S.V.** performed research, analyzed data, designed experiments and wrote the paper. **E.M.S.**, **F.W.O.** and **J.M.W.** performed experiments. **Y.B.** analyzed data. **T.A.B.** designed experiments. **T.D.** designed experiments and wrote the paper.

